# The long term effects of uncoupling interventions as a therapy for dementia in humans

**DOI:** 10.1101/2023.06.05.543670

**Authors:** Alan Holt, Adrian Davies

## Abstract

In this paper we use simulation methods to study a hypothetical uncoupling agent as a therapy for dementia. We simulate the proliferation of mitochondrial deletion mutants amongst a population of wild-type in human neurons. Mitochondria play a key role in ATP generation. Clonal expansion can lead to the wild-type being overwhelmed by deletions such that a diminished population can no longer fulfill a cell’s energy requirement, eventually leading its demise. The intention of uncoupling is to reduce the formation of deletion mutants by reducing mutation rate. However, a consequence of uncoupling is that the energy production efficacy is also reduced which in turn increases wild-type copy number in order to compensate for the energy deficit. The results of this paper showed that uncoupling reduced the severity of dementia, however, there was some increase in cognitive dysfunction pre-onset of dementia. The effectiveness of uncoupling was dependent upon the timing of intervention relative to the onset of dementia and would necessitate predicting its onset many years in advance.

## 1 Introduction

The risk of developing dementia increases with age [1] and, as a consequence of an aging population, neurodegenerative diseases and dementias have become a growing burden on society. Therefore, there is a need for interventions that can help to ameliorate cognitive decline [2]. Neuron loss is the primary cause of cognitive decline associated with dementia [3, 4, 5].

Mitochondria are generally accepted as the major source of free radicals in the cell [6, 7], albeit, there may be other sources that are as important [8]. As the mitochondrial DNA (mtDNA) is in close proximity to the electron transport chain, it will be exposed to, and damaged by, these free radicals [9]. This is the basis of the free radical theory of aging [10], which has also been proposed as a causal factor in dementia [11].

Free radical damage can cause mtDNA deletions (mtDNA_*del*_) [12], which can further impair mitochondrial function and increase free radical leakage [13]. This can lead to a vicious cycle of increasing mtDNA damage [14]. The greater the proportion of respiratory impaired mitochondria, due to mtDNA_*del*_ for example, is predictive of clinical severity such as loss of neurons [9].

We developed a simulation model based upon these reports to study mtDNA_*del*_ generation and proliferation. In this paper we investigate the long term effects of a hypothetical protonophoric uncoupling agent on a population of mtDNA within a pseudo organelle.

Clonal expansion of one or more mtDNA_*del*_ species can lead to the mtDNA wildtype (mtDNA_*wild*_) being overwhelmed, causing the cell to die. For post-mitotic cells, such as neurons, losses are permanent and will cause cognitive decline. In old age some healthy aging is to be expected but severe neuron loss can lead to dementia [3].

In previous work we used this simulator to study factors that can inflict a predisposition to dementia on an individual, namely, mtDNA half-life [15], mutation probability [16] and copy number [17]. In this paper, we focus on mutation probability and wild-type copy number (*CN*_*wild*_). In previous papers we fixed the mutation probability and per-mtDNA energy production rates (and, therefore, *CN*_*wild*_) for the entirety of a host’s lifetime. With uncoupling, we schedule changes to these factors at some point in the simulation run.

Uncoupling has the effect of reducing free radical leakage and, therefore, lowers the mutation probability. However, a consequence of uncoupling is that per-mtDNA energy production efficacy is reduced. In our simulator we implement an energy deficit/surplus feedback mechanism whereby mtDNA replication is turned on if the cell is in energy deficit and turned off if in surplus. The consequence of reducing energy production efficacy is that the deficit/surplus feedback mechanism increases the mtDNA_*wild*_ copy number to compensate for a deficit in energy.

Using our simulation, we study the potential effect of a therapeutic intervention with a hypothetical protonophoric uncoupling agent on cognitive dysfunction in humans. The simulated uncoupling agent halves mutation probability and per-mtDNA energy production efficacy. Interventions are performed at various ages, namely, 30, 50 and 65 years. We compare neuron loss with a control which receives no uncoupling therapy. Furthermore, we consider two cases: a host with low predisposition to dementia and a host with high predisposition.

Our results showed that uncoupling reduced the severity of dementia postonset. However, it also resulted in increased healthy aging pre-onset. A finding of this research is that timing of an uncoupling intervention was critical. Intervention at the onset of dementia was ineffective, whereas, if intervention was too early, there was a negative effect on a host’s cognitive function in general. Therefore, effective uncoupling therapies would require predicting a host’s onset of dementia. Furthermore, this prediction needs to be many years ahead of time which, given the state-of-the-art, is not possible.

The contribution of this paper to uncoupling as a therapy for dementia is threefold:

- Using simulation to investigate long-term effects of uncoupling on mitochondrial dysfunction.
- Demonstrating that in silico methods can overcome limitations of both in vivo and in vitro methods.
- Shows that, while uncoupling can reduce the severity of dementia, intervention needs to be performed many years prior to dementia onset and that the timing of the intervention, relative to onset, is critical.

## 2 Uncoupling

For several decades after the seminal paper by Harman [18], mitochondrial dysfunction and reactive oxygen species were thought to be a proximal cause of neurodegenerative disease [19, 20, 21]. The free radical hypothesis was superseded by the hypothesis that misfolded proteins had a central role in neurodegenerative disease [22, 23], however, in spite of decades of research there is considerable evidence that there is poor correlation between *β*-amyloid and cognitive decline [24, 25]. Furthermore, despite over 200 clinical trials the *β*-amyloid hypothesis is notable for its lack of effective therapies [26].

More recent observations indicate that cell loss is responsible for the cognitive decline seen in dementia [27]. Therefore, there has been a resurgence in the mitochondrial dysfunction and free radical hypothesis [28, 29]. There is substantial evidence that oxidative stress [30, 31], mitochondrial dysfunction [32, 33, 34, 35] and cell death due to oxidative damage [36] are critical factors in the progression of various dementias.

Antioxidant therapies that scavenge free radicals have failed to prevent the progression of dementia [37, 38]. This is to be expected given that reaction rates between free radicals and non-enzymatic antioxidants are dwarfed by the cell’s natural free radical defence system [39]. However, reducing the generation of free radicals by modulating uncoupling proteins is a potential therapeutic target for the treatment of neurodegenerative disease [40, 41, 42, 43]. This approach is supported by the observation that overexpression of uncoupling protein 2 (UCP2) in cells of the substantia nigra, in vitro, decreased free radical production; whereas, free radical production was increased in the absence of UCP2 [44].

In this paper we consider a hypothetical uncoupling agent that reduces mutation probability and (indirectly) increases *CN*_*wild*_ within the simulation environment at designated ages. Uncoupling of mitochondria can be achieved by pharmacological intervention. For example, 2,4-dinitrophenol (DNP) is a protonophoric uncoupling agent that has been recently studied for its potential neuroprotective properties.

When administered at low doses, DNP has been shown to reduce oxidative damage and increase longevity in mice [45]. It has also demonstrated potential anti-amyloidogenic and neuritogenic actions in vitro [46, 47, 48]. These in vitro studies and the in vivo experiments with mice are, by necessity, short term while the lifespan of a human neuron is in the order of decades. Nevertheless, these studies have led to a renewed interest in the use of uncoupling agents for the treatment of neurodegenerative diseases [49, 50].

One possible explanation for the beneficial effects of DNP is that it reduces the mitochondrial membrane potential (ΔΨ) leading to a decrease in the generation of free radicals [51, 52, 42, 49, 53, 54].

At low concentrations DNP effectively mimics the action of endogenous uncoupling proteins which, through proton cycling, account for 20-25% of basal metabolic rate in rats [54]. Uncoupling proteins are activated by superoxide and are neuroprotective by reducing the formation of free radicals [55]. It has been proposed that the original function of uncoupling proteins was to decrease the rate of free radical formation [56] rather than thermogenesis. A consequence of the administration of DNP (uncoupling mitochondria to reduce free radical formation) is that it could impair the generation of ATP [57, 58]. An increase in the AMP/ATP ratio will be detected by AMPK which is a conserved sensor of low intracellular ATP levels and will induce mitochondrial biogenesis [59, 60, 61].

While uncoupling could potentially decrease free radical formation and prevent neurodegenerative disease, the administration of DNP comes with risks. DNP reduces the rate of ATP synthesis and generates excess heat and, at high concentrations, could potentially lead to hyperthermia and death.

The generation of free radicals is dependent on, and sensitive to, changes in ΔΨ [53], however, this relationship is nonlinear. Below a certain ΔΨ threshold the rate of formation of free radicals is negligible; above this threshold, the rate increases exponentially. A precise value for a neural ΔΨ threshold, in vivo, is unknown but published estimates, in vitro, vary between 80 and 140mV [62, 63]. The extent to which an uncoupling agent will reduce the formation of free radicals will depend on the initial ΔΨ which, in turn, will depend on the cell type and current rate of energy expenditure.

It is known that free radicals can damage mtDNA and cause mutation but it is not known what rate of mutation will occur for a given rate of free radical generation, however, there is experimental evidence that indicates that doubling the rate of generation of superoxide will approximately double the rate of mutation [64] in vitro.

Given the difficulty of obtaining precise values for these parameters, in our model we have used values that are physiologically plausible, but are subject to revision based on future experimental determinations.

It is challenging to verify these results against any empirical data given most studies in vivo involve testing on rodents and are performed over short time periods; as are in vitro methods. The advantage of in silico methods are that they enable us gain insight into the long term effects of such therapies.

## 3 Simulator

We simulate the proliferation of mtDNA_*del*_ amongst a wild-type species within a pseudo mitochondrial organelle. We describe the simulator (written in Python) in this section which is based upon the Mitochondrial Free Radical Theory of Aging (MFRTA). In our simulator mtDNA are software abstractions that, like their biological counterparts, are subject to *expiration, replication, mutation* and *competition*.

Simulations operate in discrete time intervals *t*, where *t* is 15 minutes and each simulation is run for 100 years. At time interval *t* = 0 the simulator is initialised with an mtDNA_*wild*_ population of copy number *CN*_*wild*_ = 1, 000. In each subsequent interval *t >* 0 mtDNA can replicate (under certain conditions) and randomly mutate. Random deletion mutations lead to heteroplasmy within the organelle such that the overall mtDNA population comprises both mtDNA_*wild*_ and mtDNA_*del*_. Deletion mutations in our simulation are idealised in that, an mtDNA_*del*_ is half the size of its parent.

We have used this simulator to study the effects of half-life [16], mutation probability [15], and copy number [17] on mitochondrial dysfunction and the effect it has on neuron loss over time. In this study, we investigate uncoupling whereby we affect mutation probability and copy number of the host at some age. We provide a detailed description of the simulator below.

### 3.1 Expiration

mtDNA operate in a hostile environment and are subject to oxidative stress due to free radicals. Consequently, mtDNA expire with a half-life of 10-30 days [65, 66]. In our simulation we subject mtDNA to random damage pursuant to a 30 day halflife. mtDNA aging is simulated by assigning a time-to-live (TTL) to each mtDNA which is decremented each time interval *t* according to a Bernoulli trial success:

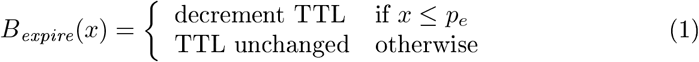

where 0 *≤ x ≤* 1 is a uniformly distributed random number. When the TTL reaches zero, the mtDNA expires. The TTL and the probability *p*_*e*_ are calibrated to yield a half-life of 30 days (see Table 3).

### 3.2 Replication

Cells need to maintain a population of mtDNA despite mtDNA having a shorter lifespan than the neurons they occupy. mtDNA replicate by cloning, thus, replenishing the population within the organelle. In each time interval *t >* 0 an mtDNA can replicate subject to certain conditions, namely:

- The cell has an energy deficit.
- The population of mtDNA does not exceed the organelle’s capacity *C*_*max*_.
- The mtDNA is not currently undergoing cloning.

The cell’s demand for energy is modelled as a consumer/producer mechanism. Each mtDNA_*wild*_ produces *e*_*wild*_ while the cell consumes *E*_*cell*_ every *t*. Thus, for each *t* there is an energy level *E*(*t*), such that:

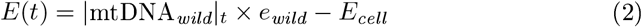

where 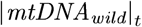 is the number of mtDNA_*wild*_ at any given interval *t*. If energy for the cell is in deficit, cloning is enabled but if there is a surplus cloning is disabled. The energy deficit/surplus mechanism maintains *CN*_*wild*_ for a given *e*_*wild*_ and *E*_*cell*_ (provided the mitochondrion is not at capacity). The cloning deficit flag CL_*def*_ is set accordingly:

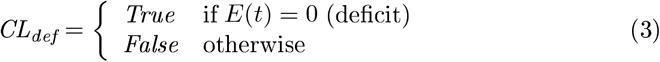

If *C*_*max*_ is the capacity of the organelle, then cloning is enabled only if |*mtDNA*|_*t*_ *≤ C*_*max*_ and is disabled otherwise. Cloning is re-enabled once |*mtDNA*|_*t*_ drops below *C*_*max*_ due to attrition of mtDNA. Thus, the cloning capacity flag 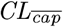 is set accordingly:

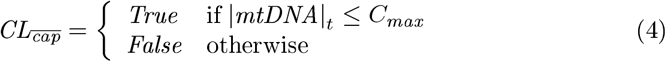

where the 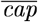 subscript means *not* exceeding capacity.

Cloning is a sequential process, that is, an mtDNA cannot simultaneously clone multiple times and must complete its current cloning process before entering the next. The time to clone is proportional to an mtDNA’s size; given that mtDNA_*del*_ are smaller than mtDNA_*wild*_, deletions have a replicative advantage over wild-type [15]. An mtDNA_*wild*_ is 16,569 nucleotides in size and takes two hours to replicate. Any mutated child is half the size of its parent and, therefore, takes half the time to replicate. For any mtDNA *m*, we compute the replication time (busy period) *T*_*busy*_(*m*) with the expression:

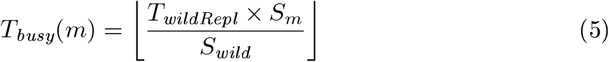

where *S*_*m*_ is the size (in nucleotides) of *m*, S_*wild*_ = 16, 569 is the size of mtDNA_*wild*_ and *T*_*wildRepl*_ = 8 time intervals (two hours). Any mtDNA *m* undergoing cloning enters a busy state for *T*_*busy*_(*m*) time intervals. The busy counter for *m* is initialised to *m*_*busy*_ = *T*_*busy*_(*m*) upon entering cloning. *m*_*busy*_ is decremented at each subsequent time interval *t* until it reaches zero.

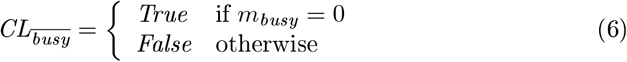

where the 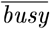 subscript means not busy.

*CL*_*def*_ and 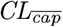 turn cloning off organelle-wide, whereas, *CL*_*busy*_ only effects a specific mtDNA. Cloning is enabled if:

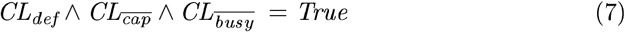

whereby a Bernoulli trial is run for the mtDNA:

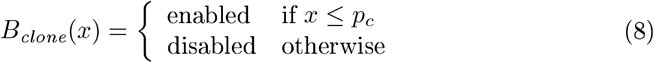

where 0 *≤ x ≤* 1 is a uniformly distributed random number. *p*_*c*_ = 0.01 and is sufficient to counter the attrition rate and maintain a population (when not in competition). A success in *B*_*clone*_(*x*) causes the mtDNA to replicate.

### 3.3 Mutation

As our model is based upon MFRTA we assume free radical damage causes deletion mutations. This is abstracted in our simulator by a Bernoulli trial with probability *p*_*m*_. Mutation takes place upon the success of the Bernoulli trial:

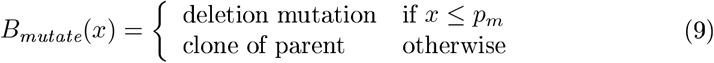

where 0 *≤ x ≤* 1 is a uniformly distributed random number.

When mtDNA undergo cloning, mutation can occur which gives rise to a new species of mtDNA. In this paper we focus solely on deletion mutations. mtDNA_*del*_ expire and replicate in the same way as mtDNA_*wild*_, thus, mutant species populations can grow in the organelle alongside the mtDNA_*wild*_. mtDNA_*del*_ can also mutate, yielding a larger deletion species.

An initial mutation probability *p*_*m*_ is set at the start of each simulation run. For the low dementia predisposition case *p*_*m*_ = 1 *×* 10^*−*4^ *h*^*−*1^ and *p*_*m*_ = 2 *×* 10^*−*4^ *h*^*−*1^ for the high predisposition case. When uncoupling is applied, the mutation probability is halved: *p*_*m*_*/*2.

### 3.4 Competition

Enforcing a maximum capacity *C*_*max*_ in the organelle creates competition for space between species. Initially, there is an abundance of space within the organelle and there is spare capacity for all species of mtDNA. However, as the mtDNA_*del*_ replicate, the aggregate mtDNA population can reach *C*_*max*_. We define the moment when the population reaches *C*_*max*_ for the first time as the competition point. At this point, space within the organelle becomes scarce and is only available after some attrition. Each species, therefore, must compete for space when cloning. When the organelle enters competition, selfish proliferation of mtDNA_*del*_ impacts the ability of mtDNA_*wild*_ to maintain energy levels within the cell. The cell dies if the mtDNA_*wild*_ population drops below a threshold (in our simulation, *CN*_*wild*_*/*3).

Figure 1 depicts a simulation run and is intended to illustrate the metrics captured from each simulation. It shows wild-type copy number (blue) and the aggregate mutant population (red). Note, when we apply uncoupling, there would be a step change in the blue line to reflect the increase in *CN*_*wild*_. Initially, the wildtype maintains a copy number pursuant to the cell’s energy needs. The green shaded area in Fig 1 shows the *competition phase*, where the left edge is the competition point *T*_*cp*_. The right edge is the point at which the cell dies *T*_*autophagy*_. The horizontal dashed line is the mtDNA_*wild*_ population threshold at which autophagy occurs. At this level the mtDNA_*wild*_ population can no longer generate sufficient energy to sustain the cell.

**Figure 1:**
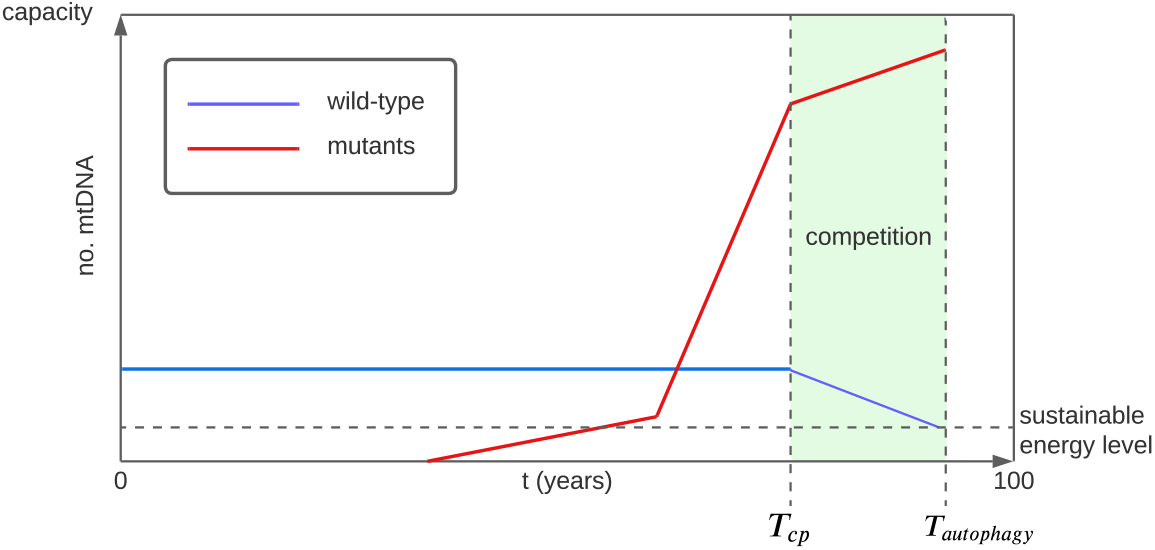
mtDNA_*del*_ proliferation

Figure 1, therefore, depicts a deceased cell, namely, one that has died before the host. A cell that never enters competition (has no competition point or autophagy edge) goes on to survive for the lifetime of the host. A cell may enter competition but may avoid autophagy and survive. In this case the competition region has a competition point edge but no autophagy edge (the right edge is 100 years, the length of the simulation).

### 3.5 Parameters

We run experiments over two factors, namely, uncoupling age (in years) and dementia predisposition (low and high). Table 1 shows the ages at which we apply uncoupling including a control simulation where no uncoupling is applied. The initial mutation probability *p*_*m*_ for the two dementia predisposition levels is shown in Table 2. Table 3 shows the fixed parameter values.

**Table 1:** The age (years) at which uncoupling intervention is introduced. CTL is the control for no uncoupling intervention.

| Designation | Age<br>(years) |
| --- | --- |
| CTL | N/A |
| U30 | 30 |
| U50 | 50 |
| U65 | 65 |

**Table 2:** Mutation probabilities associated with predisposition to dementia.

| Predisposition | Initial $p_m$<br>($h^{-1}$) |
| --- | --- |
| low | $1 \times 10^{-4}$ |
| high | $2 \times 10^{-4}$ |

**Table 3:** Simulation parameters, fixed for all simulations.

| Parameter | Value | Comment |
| --- | --- | --- |
| $t_{max}$ | 3,504,000 | Simulation run length (100 years) |
| $p_c$ | $0.01 h^{-1}$ | Cloning probability |
| $p_m$ | $1 \times 10^{-4} h^{-1}$ | Mutation probability (low) |
| $p_m$ | $2 \times 10^{-4} h^{-1}$ | Mutation probability (high) |
| $C_{max}$ | 10,000 | Organelle capacity |
| $N_{wild}$ | 1,000 | Initial mtDNA<sub>wild</sub> population |
| $T_{wildRepl}$ | 2 hours | Wild-type replication time |
| $TTL$ | 10 | Equiv. 30 day half-life |
| $p_e$ | 0.0034 | |

At the point at which uncoupling therapies are applied, the initial *p*_*m*_ and the energy production efficacy *e*_*wild*_*/*2 are halved where the consequence of halving energy efficacy is to double *CN*_*wild*_. We run 200 replications of the simulation for each factor and level. Algorithms (1) and (2) show the pseudo code for the simulator with uncoupling. Source code is available in the repository: `git clone` https://agholt@bitbucket.org/agholt/mitosim2022.git.

## 4 Analysis

In this section we show the results of uncoupling on cognitive dysfunction due to neuron loss. Cognitive dysfunction in the elderly varies from healthy aging to dementia. There is little consensus in the literature as to what cell loss rates, both global and local, constitute dementia [67, 68, 69, 27]. We use neuron loss ranges from [16, 4], that is, 20-40% loss, where the lower limit is the onset of dementia and the upper is severe. Below 20% is healthy aging and beyond 40% it is unlikely the host itself survived, however, we run the simulation for the full 100 years, regardless.

We uncouple at ages 30, 50 and 65 which are designated U30, U50 and U65 (Table 2), respectively. We also run a control simulation with no uncoupling, designated CTL.

Figure 2 shows the neuron lifespan for U30, U50 and U65 versus CTL. The left graph shows the results for a host with a low predisposition to dementia while the right shows high predisposition. We use bootstrap methods to generate sample means and plot the empirical cumulative distribution function (ECDF). For each U30, U50 and U65 the mutation probability is halved at the respective uncoupling times (*p*_*m*_*/*2) as is the per-mtDNA_*wild*_ energy production efficacy (*e*_*wild*_*/*2).

**Figure 2:**
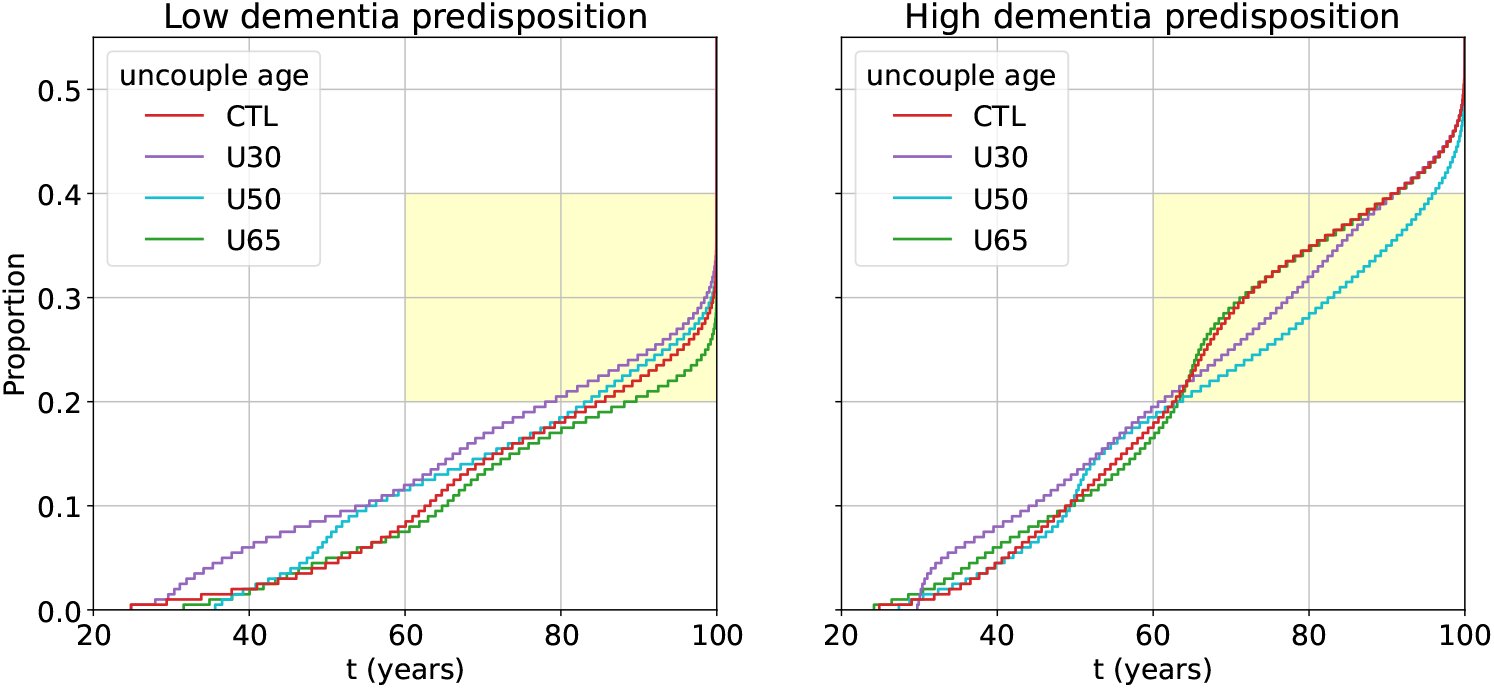
Lifespan for a host with low predispostiion to dementia (left) and a host that with high predisposition (right). The yellow shaded area represents the region of the system where the cognitive decline in the host would be classed as dementia for the elderly.

### Algorithm 1

Simulator

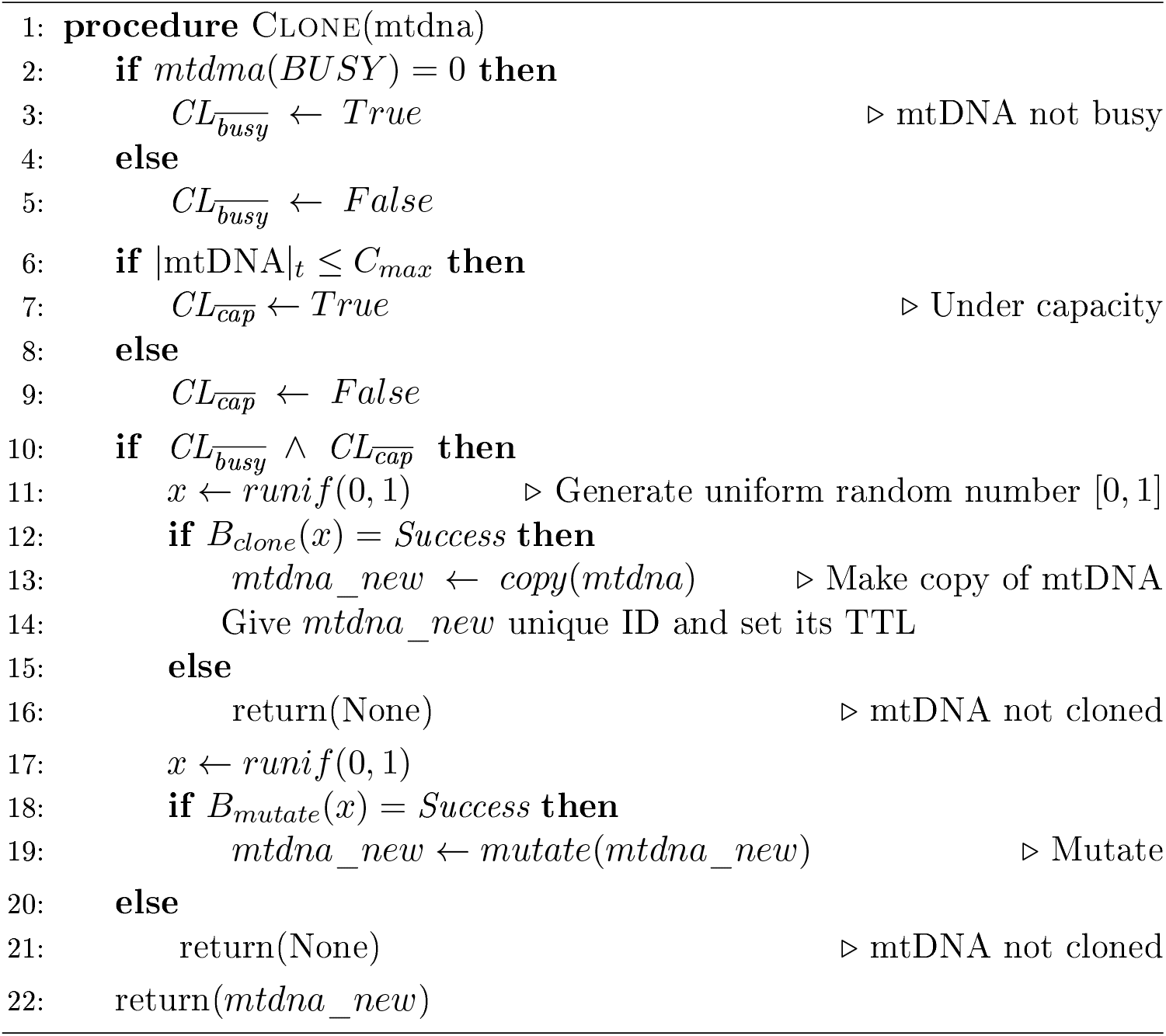

### Algorithm 2

Simulator

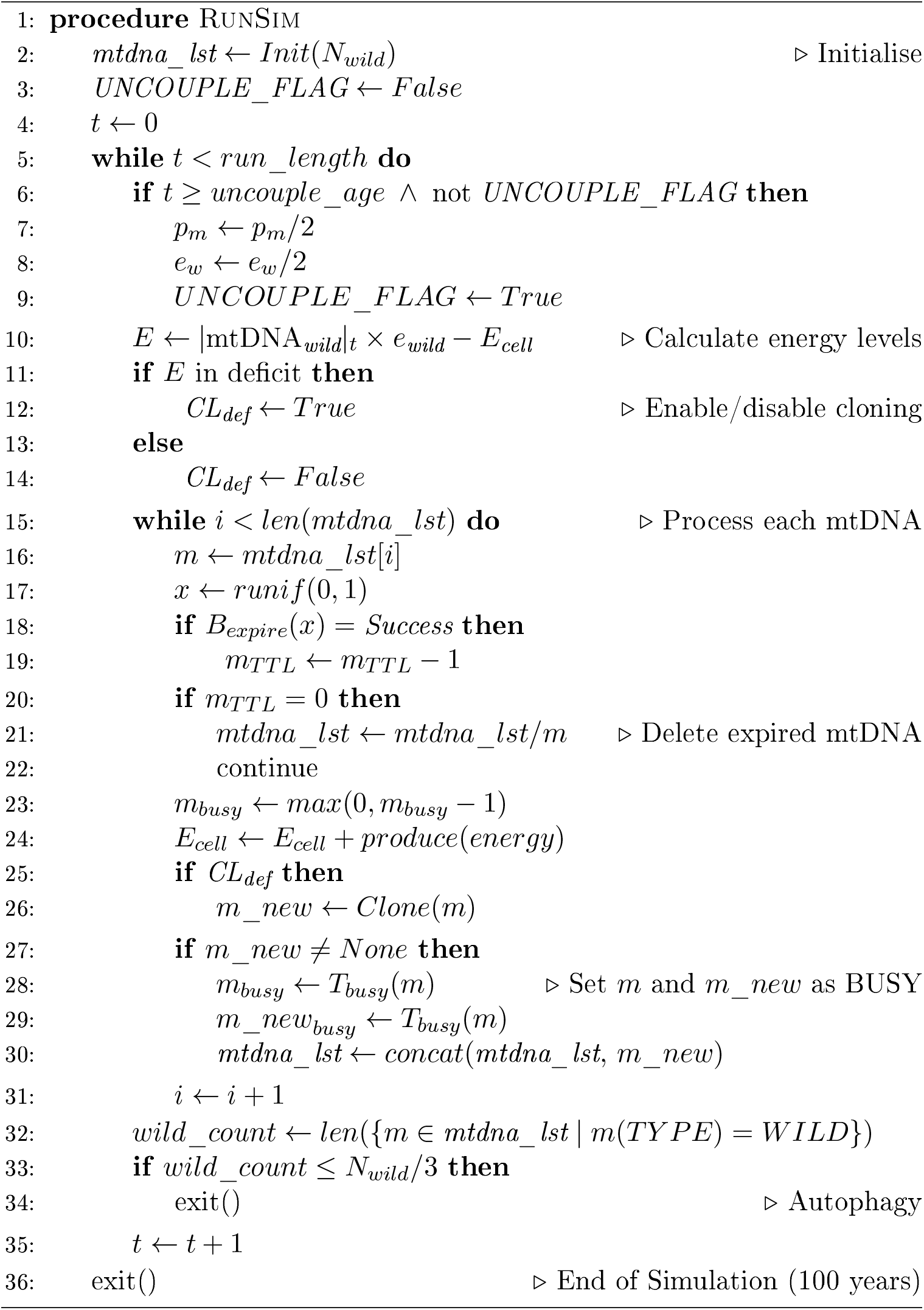

The shaded yellow area in the two graphs shows the region for dementia in the elderly (20-40% cell loss). For low predisposition, U30 and U50 compound cognitive decline compared to CTL, whereas, U65 delays the onset of dementia and reduces severity. For high predisposition, there is little difference between U65 and CTL and onset of dementia is the same regardless of uncoupling. U30 and U50 reduce the severity of dementia but increase healthy aging pre-onset. U50 yields more favourable results than U30, that is, dementia is less severe as is healthy aging.

In analysing the dynamics of mtDNA_*del*_ proliferation in an organelle we are interested in the following metrics (see Fig 1):

- Heteroplasmy of deceased cells.
- The ratio of survived cells that entered competition.
- The time at which the competition point *T*_*cp*_ is reached.
- Time in competition. For deceased cells, the time in competition is:

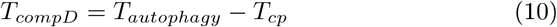

For survived cells that entered competition, the time in competition is from *T*_*cp*_ to the eventual death of the host:

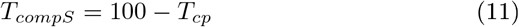

The graphs in Fig 3 shows *T*_*cp*_ (top) and *T*_*compD*_ (bottom) versus cell lifespan (for deceased cells). It can be seen that there is a strong correlation between lifespan and *T*_*cp*_ which suggests that delaying the competition point increases a cell’s chance of survival. However, correlation between lifespan and *T*_*compD*_ is not so strong.

**Figure 3:**
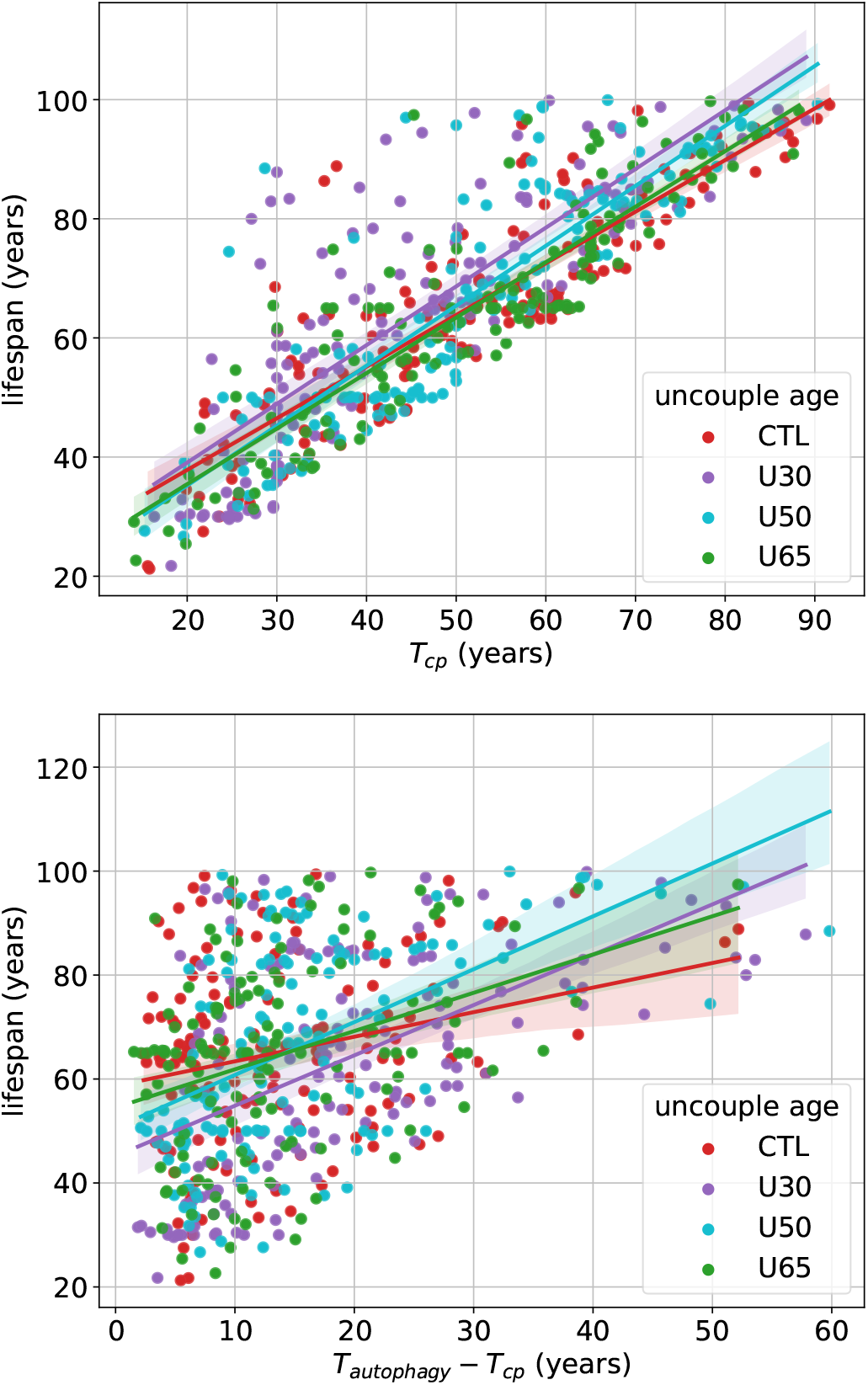
Lifespan correlated with *T*_*cp*_ and *T*_*compD*_.

Figure 4 shows deceased cell metrics for both low (left column) and high (right column) predisposition to dementia. It can be seen that, while there is a reduction in heteroplasmy (top row, Fig 4) for uncoupling compared to CTL, it is quite low.

**Figure 4:**
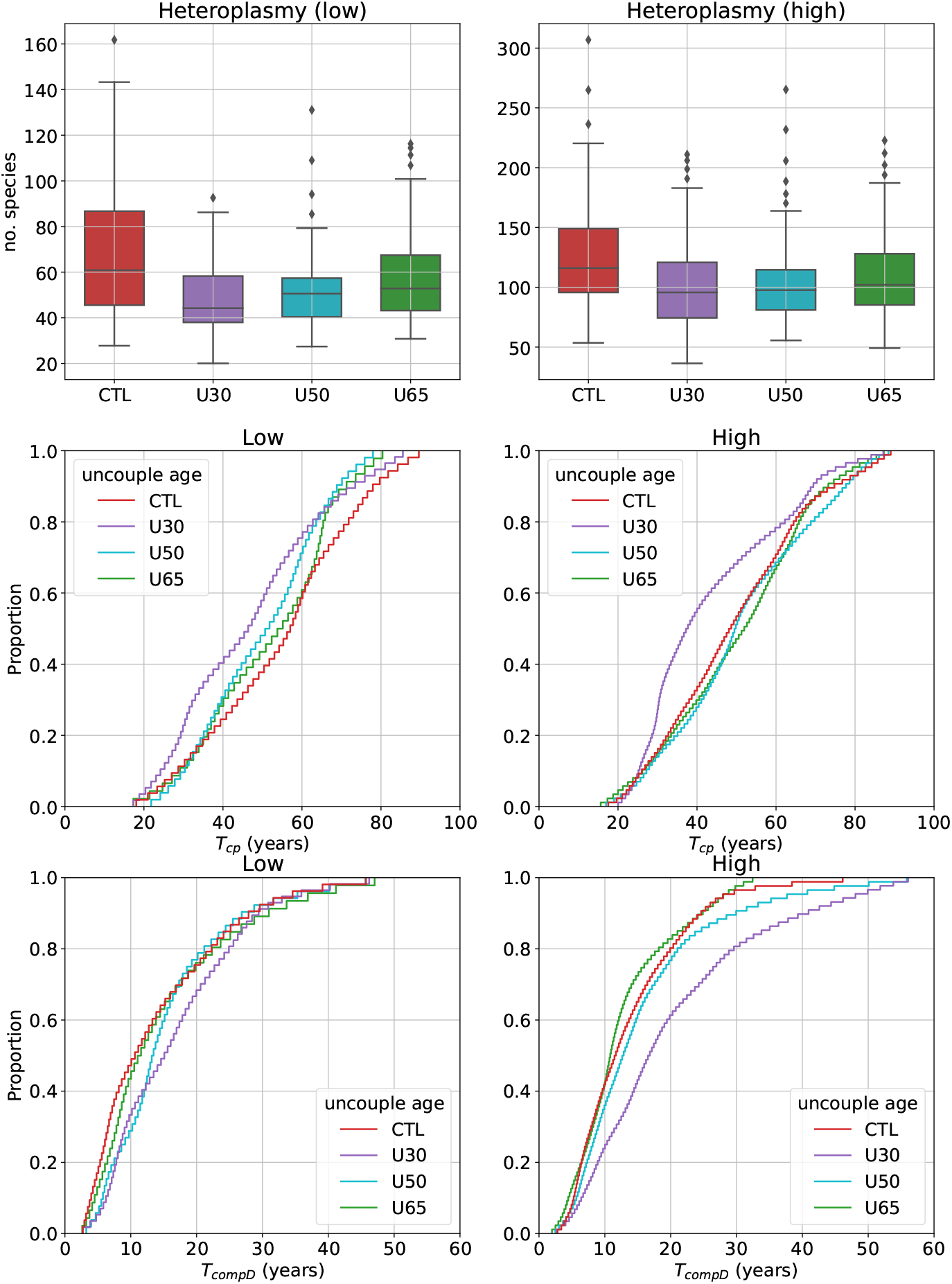
Deceased cells. Heteroplasmy is shown in the top row graphs. Competition point *T*_*cp*_ is shown in the middle row and time in competition *T*_*compD*_ on the bottom row. The left column shows low predisposition and the right, high.

The middle row shows the ECDF for *T*_*cp*_ and the bottom row shows the ECDF for *T*_*compD*_. Uncoupling brings forward *T*_*cp*_ compared to CTL. The difference for U50 and U65 is only a few years but for U30 the difference is quite pronounced. In general, uncoupling does not seem to affect time spent in competition (*T*_*compD*_) with the exception of U30 for high predisposition. If the host has a high predisposition to dementia and the intervention starts at age 30, then deceased cells spend several years longer in competition.

Figure 5 shows the metrics for survived cells that entered competition. The left and right columns are the low and high predispositions, respectively. The top graphs show the survived cells that entered competition as a proportion of all survived cells (excluding deceased cells). The proportion rises with uncoupling age, however, the chance of survival amongst cells is relatively rare.

**Figure 5:**
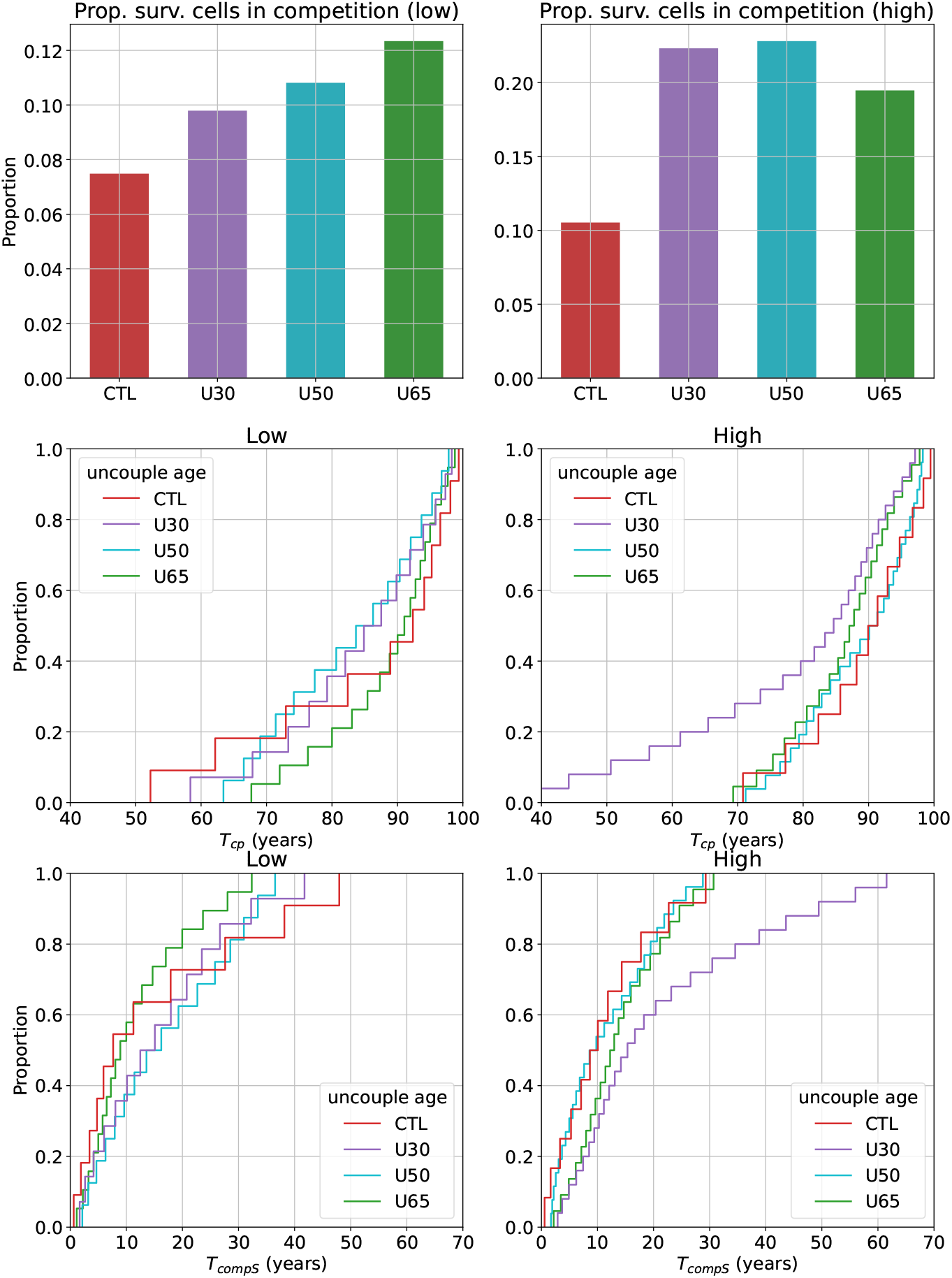
Survived cells that entered competition. The top row graphs show the proportion of survived cells that entered competition from the set of all survived cells. Competition point *T*_*cp*_ is shown in the middle row and time in competition *T*_*compD*_ on the bottow row. The left column shows low predisposition and the right, high.

The middle and bottom row graphs in Fig 5 show the ECDF for *T*_*cp*_ and *T*_*compS*_, respectively. Due to the infrequency of survived cells that entered competition, it is not easy to get reliable statistics, albeit, we generate sample means through bootstrapping. Uncoupling seems to have little effect on *T*_*cp*_ and *T*_*compS*_ with the exception of U30 for the high predisposition case where *T*_*cp*_ is significantly lower and *T*_*compS*_ significantly higher.

Figure 6 shows the heteroplasmy for survived cells that did not enter competition. It can be seen that uncoupling has little effect on this subset of cells. The data from which these graphs are derived can be found at: [Kaggle](https://www.kaggle.com/alanholt/uncoupling-analysis).

**Figure 6:**
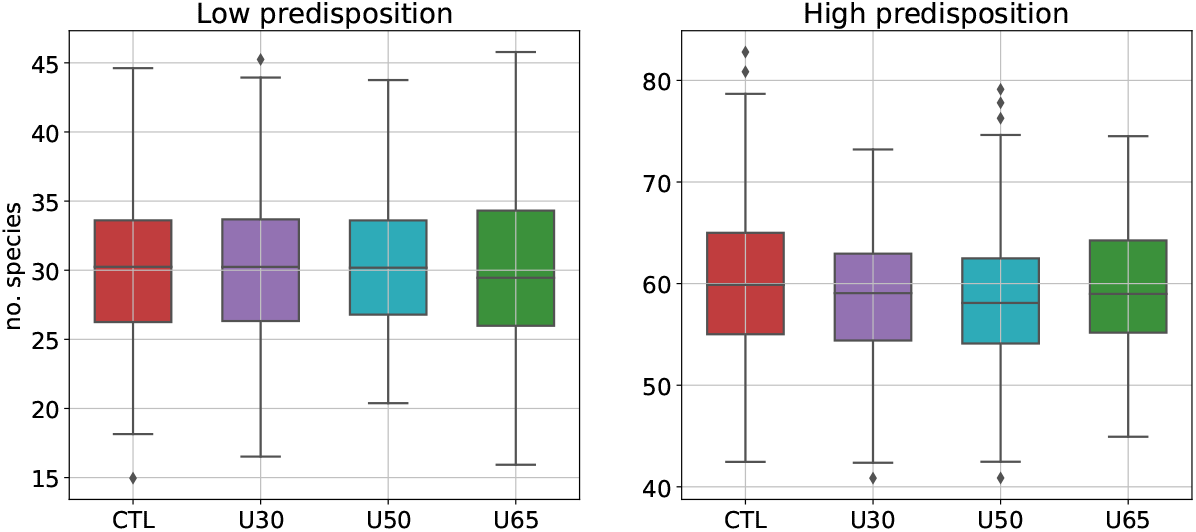
Heteroplasmy for cells that survived and did not enter competition.

## 5 Discussion

In previous studies we observed that high mutation rate and copy number were associated with a predisposition for dementia due to increased neuron loss [16, 17]. We simulated a hypothetical uncoupling agent that reduced mutation rate in the mitochondrion with a view to reducing neuron loss. Our uncoupling agent also halved per-mtDNA_*wild*_ energy production efficacy. Our simulation operates a deficit/surplus energy feedback mechanism that regulates *CN*_*wild*_, thus, the halving of energy production efficacy places the cell in energy deficit. The cell responds by turning on cloning until *CN*_*wild*_ reaches a level that can sustain its energy demands. Halving the energy production efficacy duly doubles *CN*_*wild*_.

We applied uncoupling at various ages, namely, 30, 50 and 65 years and compared the results to a control. We also considered the following two scenario, namely, low and high predisposition to dementia.

For low predisposition the onset of dementia occurred in the host’s mid-eighties for CTL. Onset was brought forward by a few years for U30 and U50 but delayed for U65. There was a small increase in dementia severity for U30 and U50 post-onset but U65 yielded a small decrease. There was a negative impact on healthy-aging for U30 and U50. We note a sudden increase in cell loss just after intervention. We attribute this to the cell suddenly finding itself in energy deficit and turning on cloning, while this increases *CN*_*wild*_ it gives species of mtDNA_*del*_ an opportunity to grow too. Thus, a host would likely experience accelerated cognitive decline in the first few years after uncoupling therapy begins.

The onset of dementia for high predisposition CTL was mid-sixties rising to severe by early nineties. The results for U65 are virtually unchanged compared to CTL. According to our analysis, a host with high predisposition will have lost approximately 50% of its deceased neurons by age 65 with approximately 30% in competition at this point. Furthermore, the host itself may already be experiencing the onset of dementia. Many of the host’s cells are in a state of clonal expansion and uncoupling necessitates an increase in the *CN*_*wild*_ due to the energy deficit of reducing the per-mtDNA_*wild*_ energy production. The cell, therefore, is trying to increase *CN*_*wild*_ in the face of stiff competition for space where mtDNA_*del*_ have a replicative advantage. The severity of dementia is reduced by U30 and U50 postonset but there is some increase in healthy-aging.

While we have considered two cases, namely, low and high dementia predisposition, human hosts would not actually be divided into these discrete categories. More realistically there would be a spectrum and the onset of dementia would vary from host to host. These results inform us that timing of any uncoupling intervention is critical relative to the onset of dementia and will vary according to where the host is on the predisposition spectrum. Interventions applied at (or beyond) the time of onset are too late as they have little effect. Interventions applied too early can have a detrimental effect on healthy aging, bring forward the onset of dementia and increase its severity.

Thus, the problem with uncoupling as a preventative therapy for dementia is predicting its onset many years ahead. As we are dealing with a hypothetical uncoupling agent we can only provide a hypothetical estimate of how far ahead an intervention needs to take place. The most effective intervention was U50 for high predisposition which would suggest intervention should begin approximately (and hypothetically) 15 years prior to onset.

## 6 Conclusions

In this paper we have used simulation to study the long term effects of uncoupling on cognitive dysfunction and dementia. In our simulation we applied a hypothetical uncoupling agent that had the effect of halving mutation probability and doubling wild-type copy number. We applied uncoupling at ages 30, 50 and 65 years and compared the results to a control that received no uncoupling therapy.

First we applied uncoupling to a host with a low initial mutation probability and thus a low predisposition to dementia. We then repeated the simulations with a higher initial mutation probability such that the host had a higher predisposition to dementia.

From these results we conclude that uncoupling can have a beneficial effect on the reduction of dementia severity. However, uncoupling can also increase healthy aging pre-onset. While we have considered two predisposition scenarios, in reality hosts would be on a spectrum with varying onsets. The benefits of uncoupling (including negative effects) appear to be dependent on when uncoupling is applied relative to onset. Interventions at or beyond the point of onset have no effect but interventions carried out too early have detrimental effects on a host’s cognition, both in terms of healthy aging and severity of dementia.

It is clear that the timing of an uncoupling intervention is critical and necessitates predicting a individual’s onset many years ahead. At the time of writing this paper there are no definitive biomarkers that can accurately predict the onset of dementia, until such time as there is, uncoupling is an impracticable therapy.

## Notes

### Competing Interest Statement

The authors have declared no competing interest.

